# A Neural Ensemble Correlation Code for Sound Category Identification

**DOI:** 10.1101/317735

**Authors:** Mina Sadeghi, Xiu Zhai, Ian H. Stevenson, Monty A. Escabí

## Abstract

Humans and other animals effortlessly identify sounds and categorize them into behaviorally relevant categories. Yet, the acoustic features and neural transformations that enable the formation of perceptual categories are largely unknown. Here we demonstrate that correlation statistics between frequency-organized cochlear sound channels are reflected in the neural ensemble activity of the auditory midbrain and that such activity, in turn, can contribute to discrimination of perceptual categories. Using multi-channel neural recordings in the auditory midbrain of unanesthetized rabbits, we first demonstrate that neuron ensemble correlations are highly structured in both time and frequency and can be decoded to distinguish sounds. Next, we develop a probabilistic framework for measuring the nonstationary spectro-temporal correlation statistics between frequency organized channels in an auditory model. In a 13-category sound identification task, classification accuracy is consistently high (>80%), improving with sound duration and plateauing at ~ 1-3 seconds, mirroring human performance trends. Nonstationary short-term correlation statistics are more informative about the sound category than the time-average correlation statistics (84% vs. 73% accuracy). When tested independently, the spectral and temporal correlations between the model outputs achieved a similar level of performance and appear to contribute equally. These results outline a plausible neural code in which correlation statistics between neuron ensembles of different frequencies can be read-out to identify and distinguish acoustic categories.

## INTRODUCTION

In the peripheral auditory system, neurons appear to encode sounds by decomposing stimuli into cardinal physical cues, such as sound pressure and frequency, which retain detailed information about the incoming sound waveform. In mid-level auditory structures, such as the inferior colliculus (IC), sounds are then further decomposed into higher-order acoustic features such as temporal and spectral sound modulations. Rather than firing when a frequency is simply present in the sound, neurons in the IC also respond selectively to spectro-temporal structure in the envelopes of frequency channels ^1^. In natural sounds, spectro-temporal modulations are highly structured and varied, and the envelopes are correlated both across frequencies and time ^2-4^. While low-level cues, such as the sound spectrum, contribute to many auditory tasks, including sound localization and pitch perception ^5,6^, spectral cues alone are insufficient for identifying most environmental sounds ^2^. Manipulating higher-order statistics related to the spectro-temporal modulations of sounds can dramatically influence sound recognition ^2^. Temporal modulations contribute to the familiarity and recognition of natural sounds such as running water sounds ^7,8^, and temporal coherence across frequencies plays a central role in auditory stream segregation ^9^. Here we examine to what extent correlated structure in the modulation envelopes of natural sounds is reflected in the correlated structure of neural ensembles and to what extent both neural and sound correlations may be useful for sound category identification. Although correlations are often thought to lead to less efficient representations of individual sensory stimuli ^10,11^, here we find evidence that correlation statistics may contribute to sound recognition and the formation of acoustic categories.

The spectro-temporal correlation structure of amplitude modulations in sounds is known to contribute to auditory perception ^2^ and strongly modulates single neuron activity in the auditory midbrain ^1^. For single neurons, correlated sound structure can improve signal detection ^12^, coding of spectro-temporal cues ^13^ and activates gain control mechanisms ^14,15^. However, how clear. Pairs of neurons at multiple levels of the auditory pathway show correlated firing that strongly depends on the spatial proximity of neurons, receptive field similarity, and behavioral state ^16-18^, but, it remains to be seen is whether these correlations are stimulus-dependent and whether they are detrimental or beneficial for coding. Theories based on efficient coding principles have proposed that stimulus-driven correlations should be minimized in order to minimize redundancies in the neural representation ^10,11^. And noise correlations, which reflect coordinated firing in an ensemble of neurons that is not directly related to the sensory stimulus, are thought to directly limit the encoding of sensory information ^19,20^. On the other hand, correlations between neurons may be functionally important and have previously been considered as plausible mechanisms for sound localization ^21^ and pitch identification ^22,23^. Here we consider a more general role for stimulus-driven ensemble correlations and test whether they may be useful for category identification and sound recognition.

The inferior colliculus receives convergent input from various brainstem nuclei and has the potential to consolidate ascending auditory information to form compact mid-level auditory representations. Here we test the hypothesis that sound correlation statistics alter the correlations between frequency organized neuron ensembles in the IC and that stimulus-driven correlation structure can directly contribute to sound recognition. First, we demonstrate that neural ensemble correlation statistics in the auditory midbrain are strongly affected by sound correlation statistics. We then show that the resulting neural ensemble statistics can be used to discriminate stimuli, and that the sound correlation structure, alone, can be used to identify and group sounds into distinct categories. In analyzing both neural and sound correlations we consider the role of spectro-temporal correlations, generally, as well as the specific contributions of temporal and spectral correlations on sound categorization. We find that stimulus-driven correlations between neuron ensembles in the auditory midbrain may provide a signature for the nervous system to recognize and categorize natural sounds.

## RESULTS

We first demonstrate that neural correlation statistics between frequency organized (tonotopic) neural ensembles in the auditory midbrain are highly structured containing both time and frequency dependent information that can be used to recognize sounds. Using a neurally-inspired sound representation, we then characterize the modulation correlation statistics of various sound categories and demonstrate how such statistics could be read out by the auditory system for sound category identification.

### Decoding Neural Ensemble Correlation Statistics to Recognize Sounds

We used multi-channel, multi-unit neural recordings in the auditory midbrain (inferior colliculus, IC) of unanesthetized rabbits to characterize and determine whether neural correlations between recording sites in the IC are affected by the correlation structure of natural sounds and whether such neural ensemble statistics could potentially contribute to sound recognition.

The spatio-temporal correlation statistics from an example penetration site demonstrate the diversity of neural ensemble correlation statistics observed in a frequency organized recording site. As expected for the principal nucleus of the IC, frequency responses areas are tonotopically organized varying from low to high frequency with recording depth (Fig. 1A; ~2-12 kHz; low frequencies more dorsal, high frequencies more ventral) and, for this reason, spatial correlations are referred to as spectral correlations in what follows. After cross-correlating the outputs of each recording site (see Materials and Methods) we find that neural ensemble correlation statistics to natural sound textures are highly diverse (Fig. 1H, I). The spectral correlation matrix (at zero-time lag) reflects the instantaneous correlation of the neural activity between different frequency organized recordings sites (Fig. 1C for fire; 1F for water) and is analogous to the frequency coincidence detection previously proposed for pitch perception ^22^. In contrast, the autocorrelograms for each recording site reflect the temporal correlation structure of the neural activity at different recording locations (Fig. 1D for Fire; 1G for water). The neural response correlation structure of the same penetration site for five natural sounds (crackling fire, bird chorus, speech babble, running water, rattling snake) are shown both along the spectral (Fig. 1H; i.e., different recording sites with different best frequencies) and temporal dimensions. We find distinct patterns for both spatial/spectral (Fig. 1H) and temporal (Fig. 1I) correlations. For instance, in this penetration site, the water and crowd have a similar spatial correlation matrices with the highest correlations localized to neighboring low frequency channels (<4 kHz, along the diagonal). Correlations are much more extensive and widespread for the fire and bird chorus, extending across the entire array. Nearby channels are still highly correlated (along the diagonal), but distant recording channels also have high correlations. The rattling snake sound, by comparison has its own distinct correlation patterns where channels with best frequencies above ~2 kHz are strongly correlated with one another.

**Figure 1:**
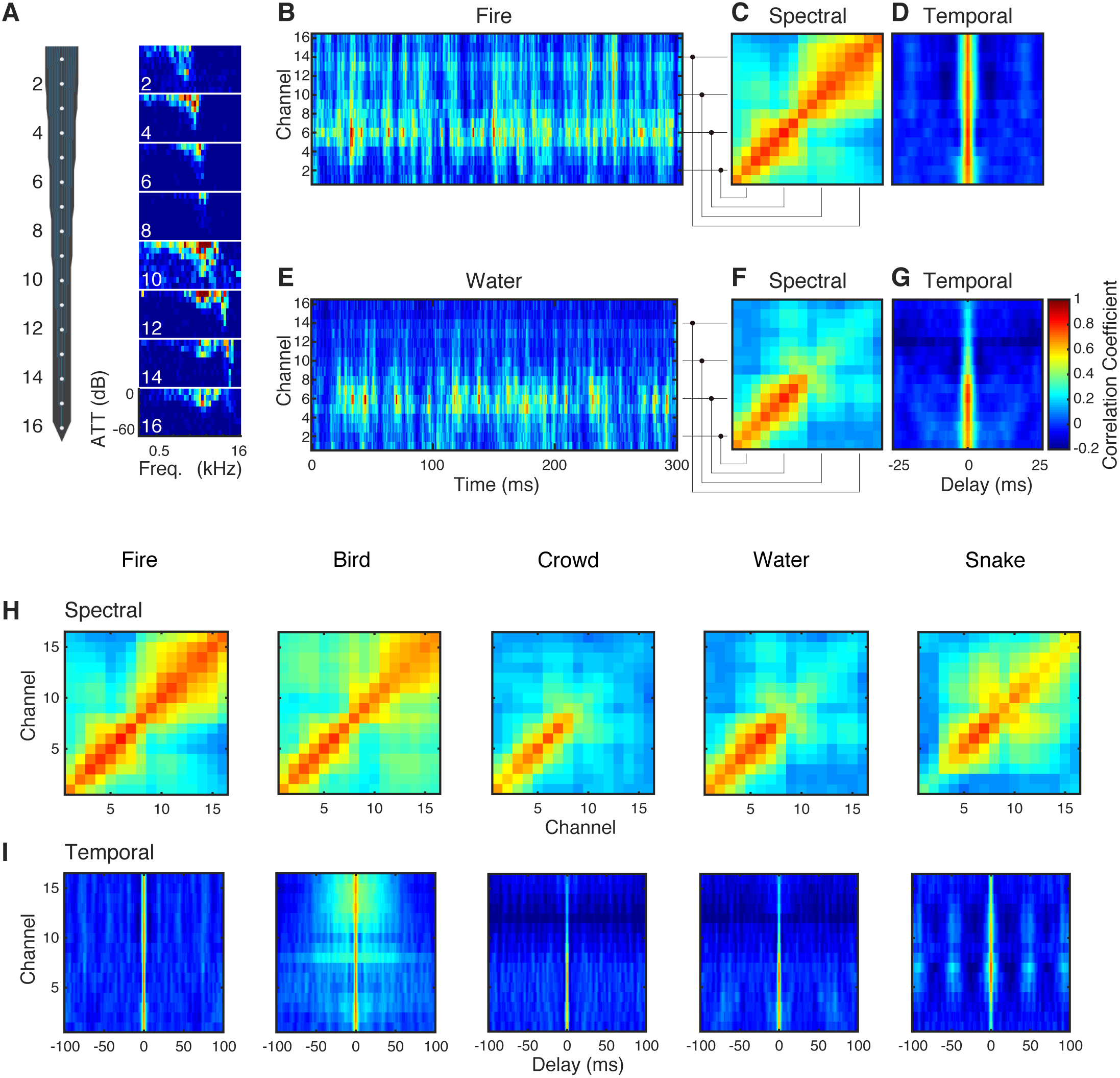
Neural ensemble correlation statistics for an auditory midbrain penetration site. (A) Neural recording probe and the corresponding frequency response areas at 8 staggered recording sites show tonotopic organization (red indicates high activity, blue indicates low activity). aMUA activity for the 16-recording channels for a (B) fire and (E) water sound segment (red indicates strong response, blue indicates weak response). The spectral (C=fire; F=water) and temporal (D=fire; G=water) neural ensemble correlation for the penetration site. (H) Spectral and (I) temporal correlations of the recording ensemble show distinct differences and unique patters across the five sounds tested.

In the time-domain, the temporal correlations of the neural activity likewise have unique patterns and time-scales for each sound (Fig. 1I). The temporal correlations for the crowd and water sounds are relatively fast (Fig. S3C; 3.0 ms, width at 50% of the maximum; 4.9 and 5.9 ms width at 10% of the maximum). By comparison, although the bird chorus sounds have a similar sharp temporal correlation (4.9 ms half width), there is also a broader and substantially slower component (112 ms, 10% width). The crackling fire produces similarly fast neural correlations (2.9 ms half width; 6.9 ms, 10% width), however these extend across the entire recording array (~1-8 kHz). This pattern likely arises because of the impulsive character of the popping ambers which are both brief in time and broadband in frequency. By comparison, although the rattling snake sound also has a precise temporal correlation at zero lag (3.9 ms, 50% width; 8.9 ms, 10% width) the ensemble generates a broad periodic pattern with a period of ~50 ms reflecting the structure of the rattling at ~20 Hz.

Since the structure of the neural ensemble correlations is highly diverse, we next quantify to what extent these statistics could be used to identify sounds (Fig. 2). We use a cross-validated minimum distance classifier (see Materials and Methods) to determine whether the spectral or temporal neural correlation structure could distinguish amongst the five sounds delivered. Neural classifier performance results for spectral, temporal, and spectro-temporal classifiers are shown for the penetration site shown in Fig. 1 as a function of sound duration (Fig. 2A). For short durations (62.5 ms), the spectro-temporal classifier performance for each sound is quite variable, although the average performance is above chance level (20%). As expected, the individual sound and average classifier performance improves with the duration of the recording. When we consider spectral correlations in isolation, the spectral classifier performance is nearly identical to the full classifier (spectro-temporal) and has similar trends for each of the individual sounds. By comparison, the temporal classifier has somewhat poorer performance.

**Figure 2:**
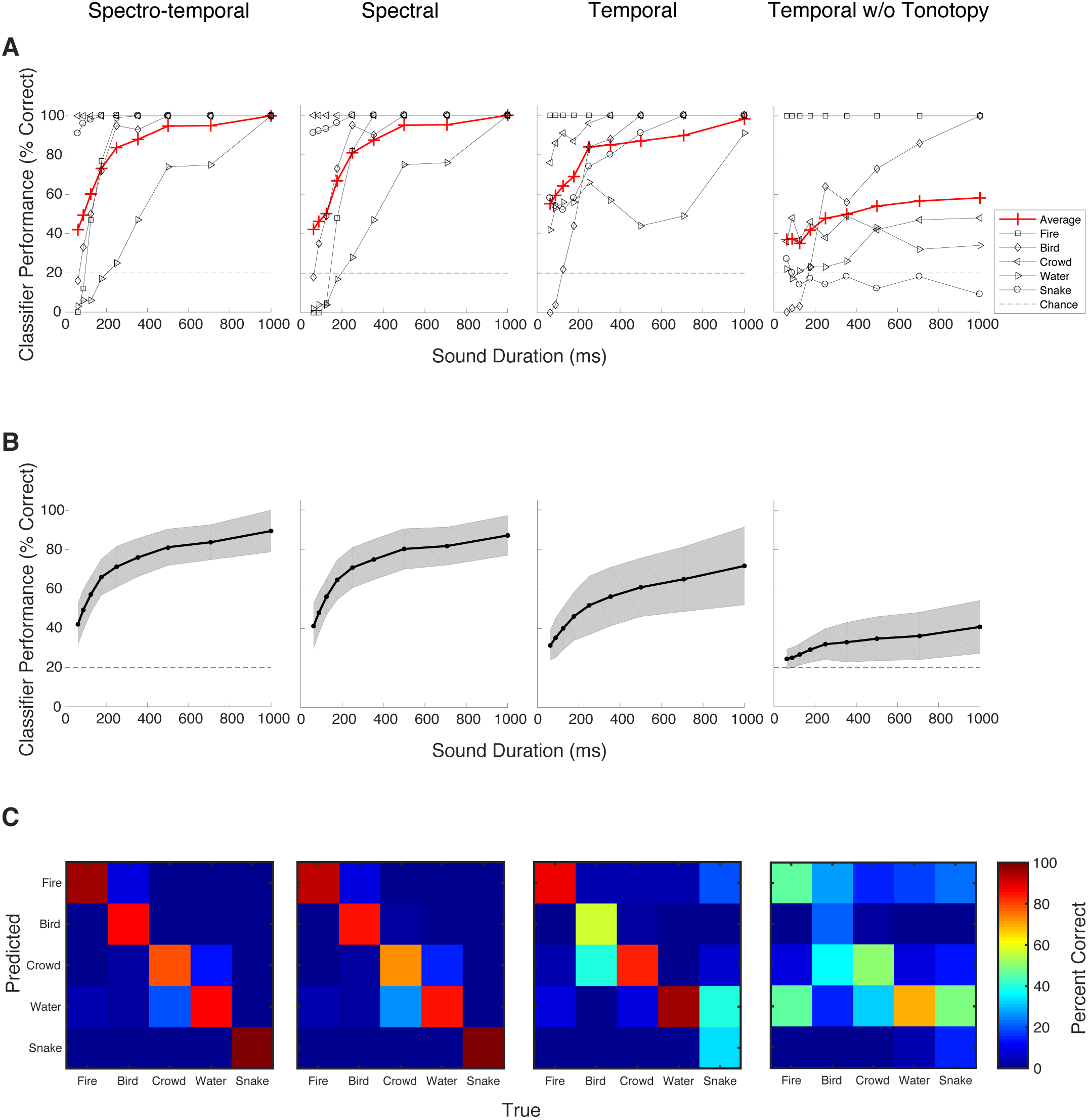
Using neural ensemble correlation statistics to identify sounds. (A) Classification results for the penetration site shown in Fig. 1. The average classifier performance (red curve) and the performance for each individual sound (gray lines) are shown as a function of the sound duration for four classifiers. In all cases, classifier performance improves with sound duration. The combined spectro-temporal classifier has the highest performance, followed by the spectral and temporal classifier. Removing the tonotopic ordering of recording sites for the temporal classifier (far right) substantially reduces its performance. (B) Average performance across all of the IC penetration sites shown as a function of sound duration. Gray bands represent SD. (C) Average confusion matrices obtained across all penetration sites for the corresponding conditions shown in (B) for a sound duration of 1 second.

Although similar average trends are observed for both the temporal and spectral neural correlation, differences are observed between the performance for different sounds. For example, the fire sound has 100% classification accuracy for the temporal classifier regardless of temporal resolution used while the spectral classifier accuracy is near 0% at 62.5 ms duration and quickly transitions to 100% at ~200 ms duration (Fig. 2A). In general, we observe similar differences across the population where some sounds are better classified with temporal correlations compared to spectral correlations or vice versa (Fig. S1). For example, the average population performance of the temporal correlation classifier for snake remained near chance level regardless of the neural response duration (Fig. S1C) while the spectral correlation classifier performance is comparatively high and improved with increasing response duration (Fig. S1B). Examination of the neural correlations and the acoustic correlations in the rattling snake sound reveals that the periodicity of the snake rattle for the training (first half) and validation (second half) data has different modulation frequencies (~16.5 Hz vs. 20 Hz) which leads to misclassification for a number of penetration sites. By comparison, the spectral correlation structure for this same sound is stable and sufficiently unique to allow high classification performance.

While the temporal classifier performance for the example penetration site is relatively high, it is possible that frequency-dependent information contributes to the temporal classifier performance. The temporal correlation matrix used for the classification maintains the tonotopic ordering of the recording channels, and this frequency-dependence could be used to improve classification. Altering tonotopy by manipulating the frequency ordering of sounds, for instance, has been shown to impair pitch perception and vowel recognition ^6,24^. To remove the frequency dependence and measure the contribution of purely temporal correlations, we distorted the tonotopic ordering from the temporal correlation matrix. Here we randomize the ordering of the recording channels during the classification. Doing so substantially reduces the classifier performance from nearly 100% average performance to ~50% for the longest sound duration measured (1 s; Fig. 2A). Thus, removing the tonotopic ordering to isolate purely temporal correlations leads to a substantial reduction in the classifier performance.

Averaging across all penetration sites (N=13) we find that, just as with the single example, neural correlation structure is highly informative and can be used as a neural response feature to recognize sounds. Across all sounds and classifiers, the average neural classifier performance improves with increasing sound duration (Fig. 2B). As with the example penetration site (Fig. 2A), the spectral and temporal classifier performance can be very different for different sounds (Fig. S1). Distinct differences in the overall classification accuracy and confusions are also observed in the average confusion matrices (Fig. 2C, average across all penetration sites for 1 s duration sound). Furthermore, when the tonotopic organization is removed from the temporal classifier the classifier accuracy is substantially reduced (Fig. 2B, far right). The average performance trends of the spectro-temporal classifier differs substantially from the temporal classifier but followed a nearly identical trend across sounds as the spectral classifier (Fig. S1, compare A, B and C). This indicates that the performance of the spectro-temporal classifier is dominated by the spectral correlations. The performance of the temporal classifier (Fig. 2B), by comparison, is somewhat lower than the spectral classifier and removing the tonotopic organization to isolate strictly temporal correlations further reduces the classifier accuracy (Fig. 2B, far right). These differences suggest that spectral and temporal correlation statistics can contribute differentially to sound identification and that, when both are combined, spectral correlations dominate the classification performance. Overall, we find that spectral and temporal correlations in IC neural ensembles are highly structured and could serve as features for sound recognition.

### Correlation Statistics of Natural and Man-Made Sound Categories

After establishing that neuron ensembles in IC have highly structured spatio-temporal correlation statistics, we now aim to determine to what extent these correlations are inherited from the correlations in the sounds themselves. Here we use an auditory model to characterize the structure of spectro-temporal correlations in an assortment of natural sound categories and develop a Bayesian classifier to determine their potential contribution towards sound category identification. Natural sounds representing 13 acoustic categories along with white noise (as a reference) are analyzed with the auditory model. We first describe the time-averaged correlations to identify structural differences between sound categories. We then analyze short-term, time-varying correlations to characterize the temporal dynamics and nonstationarity of different sound categories.

#### Average Correlation Statistics and Diversity

Here we evaluate correlations between modulations of different frequency-selective outputs of a cochlear model (see Materials and Methods.). The time-averaged spectro-temporal correlations (Fig. 3B and F) highlight distinct acoustic differences between sounds. At zero-lag, the spectro-temporal correlations reflect the instantaneous correlations between different frequency channels (Fig. 3C and G), while correlations within the same frequency channels at different lags reflect the autocorrelations (Fig. 3D and H). For example, the spectral correlation structure of speech is relatively broad reflecting the strong commodulation between frequency channels. This contrasts the running water excerpt, where the correlations are largely diagonalized, indicating minimal correlation between distant frequency channels. The temporal correlation structure in speech exhibits a relatively slow temporal structure (64 ms half width), which reflects the relatively slow time-varying structure of speech elements and words ^25,26^. By comparison, the water sound has a relatively fast temporal correlation structure (4 ms half width) that is indicative of substantially faster temporal fluctuations in the sound power.

**Figure 3:**
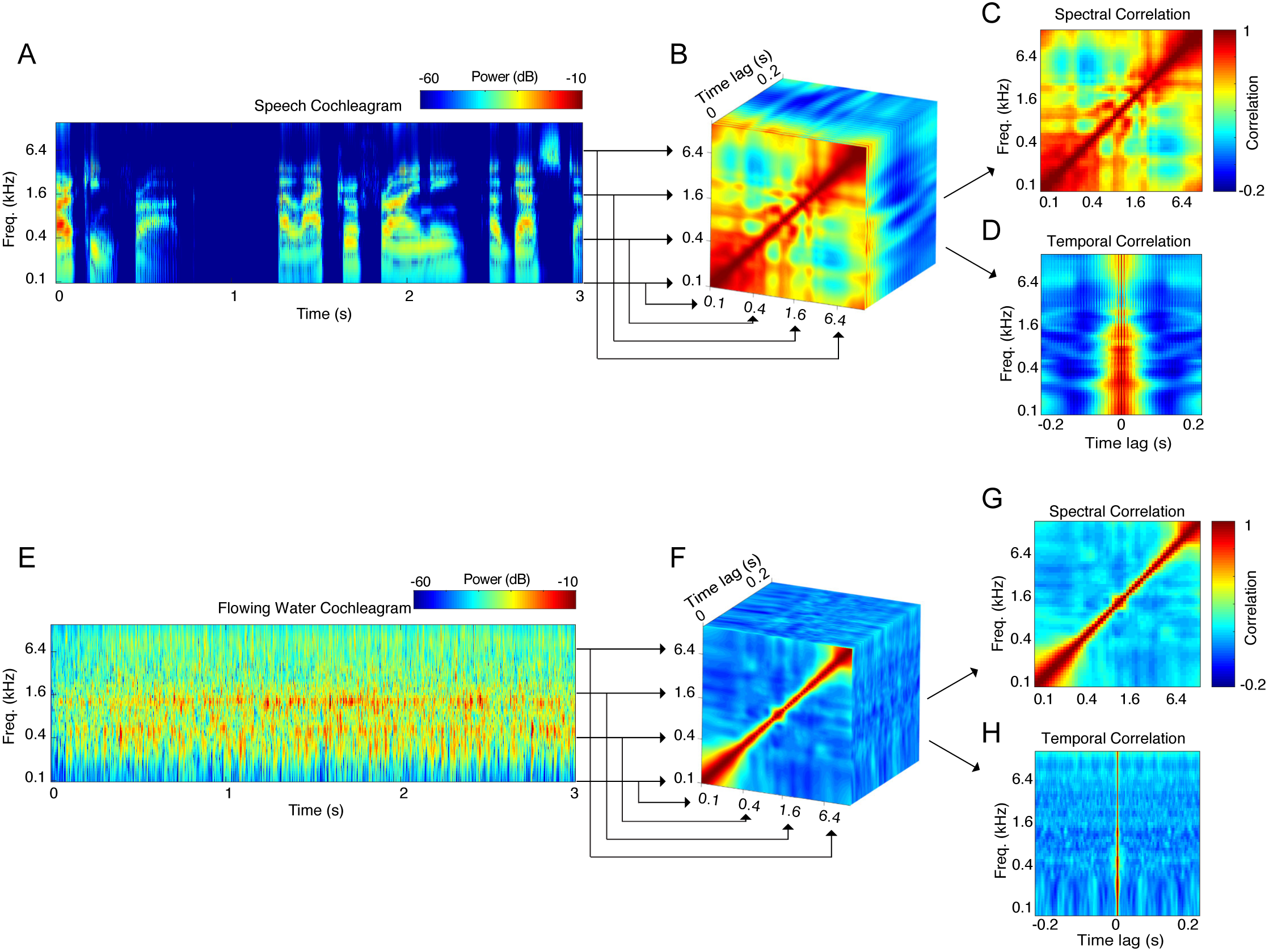
Measuring the average correlation structure of natural sounds. Illustrated for a speech and a flowing water sound. The spectro-temporal correlations are obtained by cross-correlating the frequency organized outputs of a cochlear model representation (A, E). The resulting spectro-temporal correlation matrices (B, F) characterize the correlations between frequency channels at different time-lags. The spectro-temporal correlations are then decomposed into purely spectral (C, G) or temporal (D, H) correlations. Speech is substantially more correlated across frequency channels and its temporal correlation structure is substantially slower than for the water sound.

The average spectral (Fig. 4A) and temporal (Fig. 4B) correlations of 13 sound categories and white noise reflect conserved acoustic structure in each of the categories, and, for the categories tested here, this structure is highly diverse. Certain sounds categories, particularly background sounds such as those from water (rain, running water, waves) and wind, have relatively restricted spectral correlations (diagonalized) and relatively fast temporal correlation structure (impulsive; correlation half width: water=3.9 ms; wind=3.6 ms). Such diagonalized and fast temporal structure reflects the fact that these sounds are relatively independent across frequency channels and time. Note that due to the bandwidth of the overlapping filters in the cochleogram, white noise has similar, restricted spectral correlations, rather than perfectly uncorrelated frequency channels. Other sounds, such as isolated vocalizations (e.g., cat, dogs and speech) have more varied and extensive spectral correlation that are indicative of strong coherent fluctuations between frequency channels. Such sounds also have relatively slow temporal correlation structure (correlation half width: speech=64.9 ms; dogs=76.3 ms; cats=117.3 ms), indicating slow dynamics associated with the production of vocalizations.

**Figure 4:**
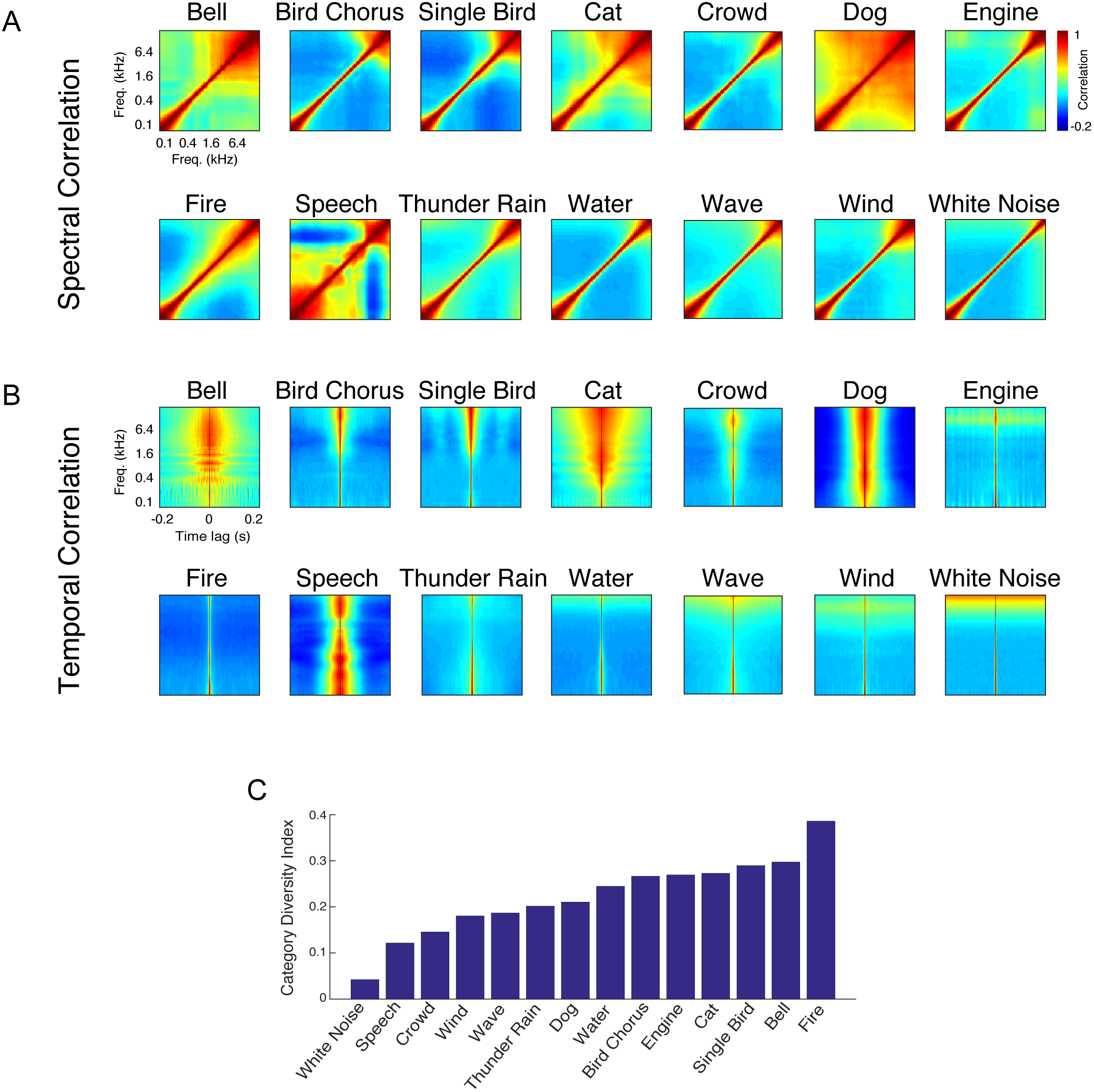
Sound correlation statistics for the thirteen sound categories and white noise. The category average (A) spectral correlation matrix and (B) temporal correlations show unique differences amongst the thirteen sounds examined. (C) The category diversity index (CDI) quantifies the variability of the correlation statistics for each category. A CDI of 1 indicates that the sound category is diverse (the correlation statistics are highly variable between sounds) while 0 indicates that the category is homogenous (all sounds have identical correlation statistics).

Although the average statistics illustrate differences between categories that could facilitate recognition, each particular sound in a given category can be statistically quite different from the average. This diversity could potentially limit the usefulness of such statistics in recognition. Sound categories in which the correlation statistics exhibit little diversity from sound-to-sound may be easier to identify, using a template-based classifier, for instance, while sounds with large amount of diversity might be expected to be more difficult to identify. For this reason, we developed a category diversity index (CDI) to quantify the diversity in the correlation structure within each of the sound categories (Fig 4C). Of all sounds in our database, white noise has the smallest diversity, which is expected given that white noise is wide-sense stationary. Of the natural sounds, speech has the lowest diversity (CDI=0.12). Although this is unexpected, it likely reflects the fact that the sound segments used in our database consisted of different speech excerpts from a single male speaker. More generally, however, there is no clear distinction or trend between different classes of sounds. For instance, the diversity indices for isolated vocalization categories are quite varied. Bird songs have a relatively high CDI of 0.29 and barking dogs had intermediate values (0.21). Similarly, the diversity of background sounds is quite varied ranging from highly diverse categories such as fire (CDI=0.39) to less diverse categories such as crowd noise (CDI=0.15) and wind (CDI=0.18). Although such trends partly reflect biases in the selection of sounds in the database, they also likely reflect acoustic properties that are unique to each sound category.

#### Short-Term Correlation Statistics and Stationarity

Although the average correlation statistics provide some insights into the large-scale structural differences between sound categories, many sounds, such as vocalizations, exhibit nonstationary structure with complex temporal dynamics and are not well described by time-averaged correlations. Here, to characterize this nonstationary structure, we use time-varying, modified short-term correlations ^27^. Computations involving temporally localized and continuously-varying correlations may be more plausible than time-averaged correlations, since lemniscal auditory neurons integrate sounds over restricted integration time windows (approximately 10 ms in the midbrain to 100 ms in cortex ^18^).

The short-term correlation decomposition of a speech excerpt and a flowing water sound are shown in Fig. 5 (A-D for speech; E-H for water; see supplementary movies, M1-M4). For these examples, we compute the time-varying correlation functions of the cochleograms using a 400 ms moving window, and these two examples illustrate the extreme differences that are possible with the time-varying statistics. Speech is highly nonstationary at this time-scale and the correlation structure (spectro-temporal, spectral and temporal correlations) varies considerably from one instant in time to the next. Such nonstationary spectro-temporal correlation structure is, in part, due to the differences between periods of speech and silence. However, even within speech periods the correlation structure can vary and can be quite distinct for each word. These differences likely reflect the range of articulatory mechanisms involved during speech production and the rapidly changing phonemes, formants, and pitch. In sharp contrast to speech, water sounds are relatively stationary. For an example sound, the correlation statistics at each instant in time are relatively consistent and closely resemble the time-averaged correlation (Fig. 5B-D and F-H; average shown in gray boundary). No matter which time segment we examine, spectral correlations are diagonalized indicating that only neighboring channels have similar envelopes, while temporal correlations are relatively fast with similar fast time constants across all frequencies.

**Figure 5:**
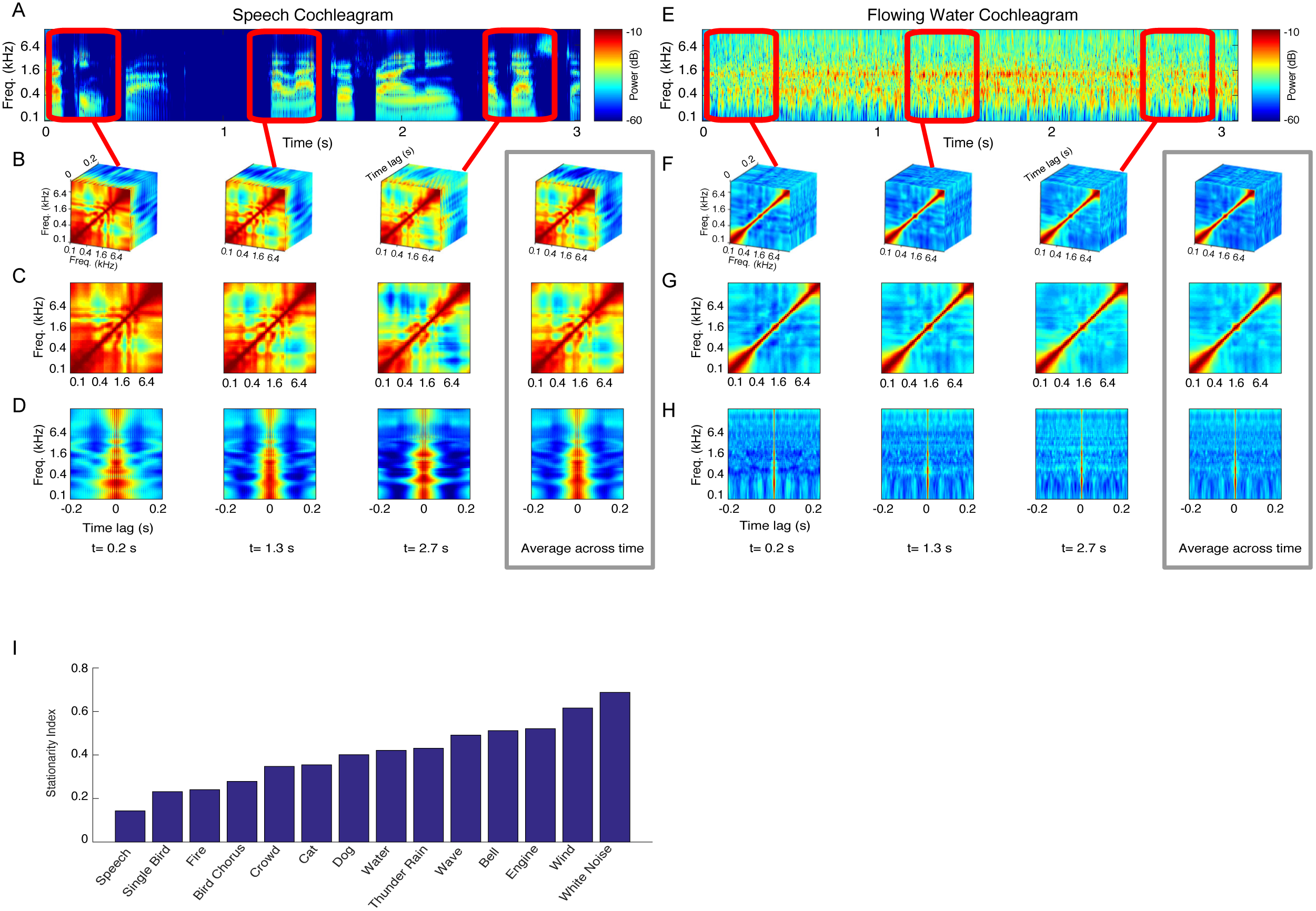
Short-term correlation statistics and stationarity. The short-term correlation statistics are estimated by computing the spectro-temporal correlation matrix using a moving sliding window. The procedure is shown for an excerpt of (A) speech and (B) water (additional examples in supplementary movies, M1-M4). The sliding window (400 ms for these example) is varied continuously over all time points but is shown for three select time points for this example. The short-term statistics are also shown for the spectral and temporal correlation decompositions. Note that for speech, the correlations change dynamically from moment-to-moment and differ from the time-average correlations (gray panel) indicating nonstationary structure. By comparison, the time-varying correlations for water resemble the time-average correlations (gray panel) indicating more stationarity. (C) Stationarity indices for the thirteen categories and white noise. Speech has lowest stationarity values while white noise is the most stationary sound.

To quantify the degree of stationarity (or lack of) in the short-term correlation function we developed a stationarity index (SI, see Materials and Methods). The SI quantifies the degree to which the sound correlation function varies dynamically over time with a value of 1 indicating perfect stationarity and a value of 0 indicating that the sound is highly nonstationary. Although the results are quite varied, we note several trends. First, except for fire sounds, environmental sounds (including running water, waves, thunder and wind) tend to have the highest average SI (SI=0.49 ±; mean ±SEM). This might be expected given that such environmental sounds typically consist of mixtures of randomly arriving sound elements (e.g., water droplets, air bubbles, etc.) and have been shown to be perceptually well described by average statistics ^2,7^. By comparison, vocalized sounds have a somewhat lower nonstationary index (SI=0.28 ±0.04; mean ±SEM) at the analysis time-scales employed. This is expected given that vocalizations transition from periods of silence and vocalizations at time-scales of just a few Hz ^26^ and even within vocalization segments the correlation structure can vary dynamically from moment-to-moment (e.g., Fig. 5). In the classification analysis that follows, we aim to describe this nonstationary structure and determine to what extent ignoring nonstationary impairs sound categorization.

### Decoding Correlation Statistics to Categorize Sounds

Given that the neural ensemble correlation statistics are strongly modulated by the correlation structure in sounds and that natural sound ensembles have highly varied spectro-temporal correlation statistics, we next tested whether these high-order correlation statistics could directly contribute to sound category identification. Here we aim to quantify the specific contributions of spectral and temporal correlations as well as the overall accuracy of classification when both features are used. While both spectral and temporal envelope cues can contribute to a variety of perceptual phenomena they often do so differentially. For instance, speech recognition can be performed with low spectral resolution so long as detailed temporal information is preserved in each frequency channel ^28^. By comparison, music perception requires much finer spectral envelope resolution ^29^. Thus, it is plausible that one of the two dimensions (temporal or spectral) could be more informative for specific sound categories or for the sound category identification task as a whole.

Rather than using a minimum distance classifier on the time-averaged correlations (i.e., Fig. 3 and 4), such as the one we used for classifying neural correlations, here we develop a probabilistic model of the short-term correlation statistics. After computing the time-varying correlations for purely spectral, temporal, or spectro-temporal correlations (Fig. 5), we reduce the dimensionality of the features using principal component analysis (Fig. S4). We then fit the low-dimensional representation of the correlations using an axis-aligned Gaussian mixture model for each sound category (see Materials and Methods). After training, we classify test sounds by comparing the posterior probability of the sounds under the mixture models of each category. By limiting the short-term correlations to spectral (Fig. 5C and G), temporal (Fig. 5D and H), or spectro-temporal (Fig. 5B and F) we can measure how each of these acoustic dimensions contributes to categorizing sounds (see Materials and Methods). Ignoring the nonstationary structure by averaging the statistics over time impaired the sound category performance, particularly for short duration sounds where the nonstationary classifiers had ~25% higher accuracy compared to the classifier based on averages (Fig. S5). The time-varying statistical structure of the sounds can, thus, contribute to more accurate sound categorization.

We optimized the model and classifier for each task separately using multiple temporal resolutions (τ_*W*_ =25-566 ms; Fig. 6A) while using the maximum sound duration (10 s). The optimal resolution for both the temporal and spectral correlation classifiers is 141 ms, while the optimal resolution for the joint spectro-temporal correlation classifier is slightly faster (100 ms). For both spectral, temporal and spectro-temporal correlations, the classifier performance is above chance for all temporal resolutions tested (Fig. 6A; chance performance = 7.69%; p<0.01, t-test, Bonferroni correction). For the spectro-temporal classifier, performance improves with increasing sound duration reaching a maximum accuracy of 84% (Fig. 6B). Individually, spectral and temporal correlations alone achieve similar maximum performance (spectral=83%, temporal=81%; p=0.60, two-sample t-test) and neither is significantly different from the overall spectro-temporal performance (spectral versus spectro-temporal: p=0.79, two-sample t-test; temporal versus spectro-temporal: p=0.42, two-sample t-test). This indicates that that both spectral and temporal correlations contribute roughly equally to the sound identification task for the full sound duration. However, the spectral classifier performance increases at a faster rate than the temporal classifier (reaching 90% of maximum in 1.7 vs. 3.0 s; Fig. 6B), indicating that evidence about the sound category may be accumulated more efficiently using spectral correlations. The joint spectro-temporal classifier improves at an even faster rate (reaching 90% of its maximum in 1.2 s).

**Figure 6:**
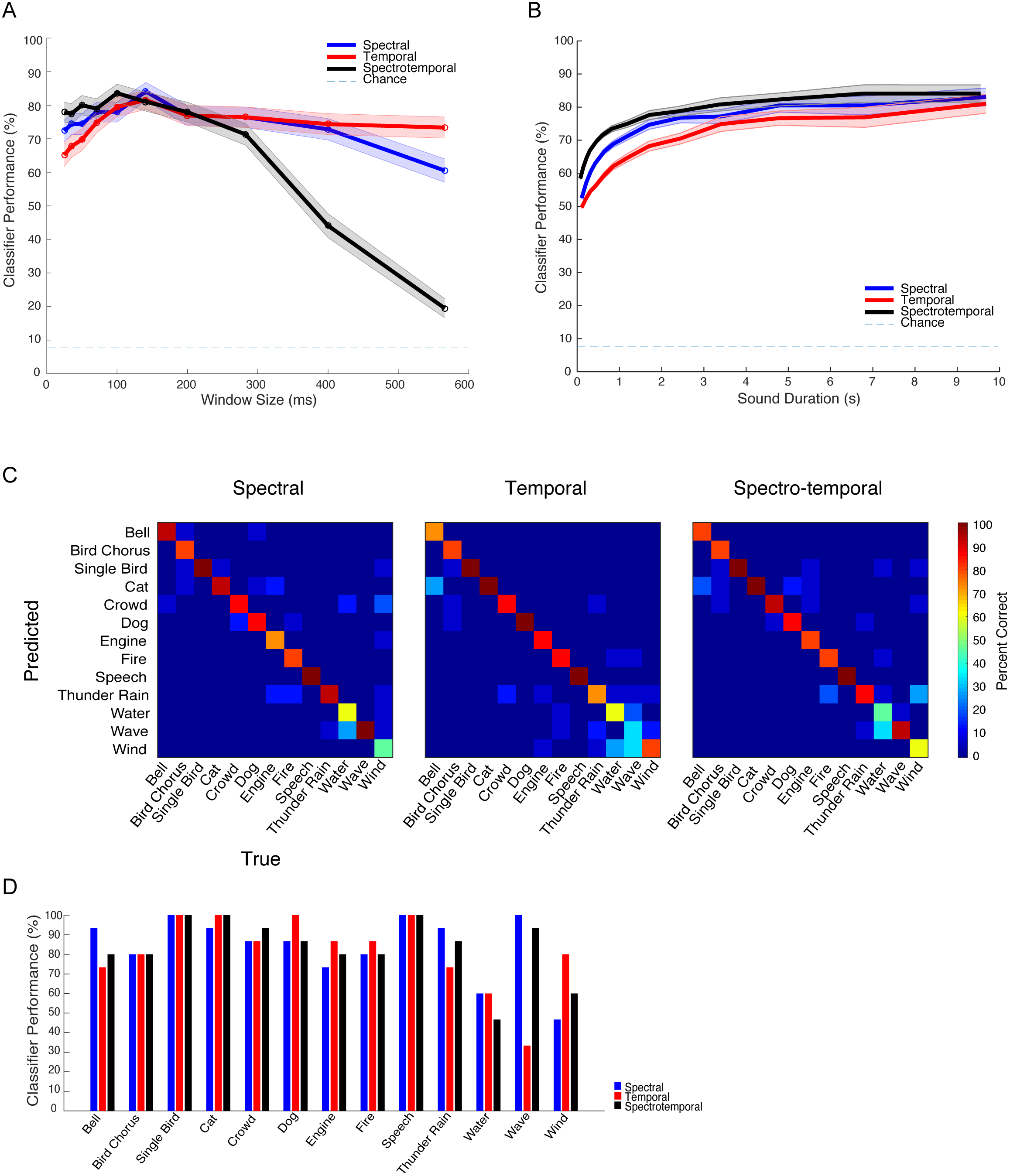
Using short-term correlation statistics to categorize sounds in a 13-category identification task. A cross-validated Bayesian classifier is applied to the short-term correlation statistics (spectral, temporal and spectro-temporal) to identify the category of each of the test sounds (see Materials and Methods). (A) Both the spectral and temporal classifiers had an optimal temporal resolution of 144 ms (i.e., short-term analysis window size). The optimal resolution of the spectro-temporal classifier, by comparison is slightly higher (100 ms). (B) For all three classifiers, the performance improves with the sound duration. The spectro-temporal classifier performance improved with sound duration at the fastest rate, while temporal correlations had the slowest rate of improvement. (C) Confusion matrices for the three classifiers for 10 s sound durations. (D) Performance for the three classifiers shown as a function of sound category (measured at the optimal resolution and at 10 s sound duration).

As with the neural correlations, we find distinct differences between the performance of the spectral and temporal classifiers for the individual sound categories (Fig. 6C and D). The recognition accuracy for each sound and the types of confusions that occur are quite different for the spectral and temporal classifiers (Fig 6C). Certain sounds such as waves perform substantially better for the spectral correlation classifier (spectral=100% vs temporal=33%). Other sounds, such as wind exhibit higher performance with the temporal classifier (spectral=47% vs. temporal=80%). Thus, although on average the performance of the spectral and temporal correlations classifier is comparable, performance of certain sound categories appears to be dominated by one of the two sets of features (Fig 6D).

We also evaluate the performance of the classifier in a two-alternative forced choice task where we require the classifier to distinguish vocalization and background sound categories. The model performance is consistently high with accuracy rates for the spectro-temporal classifier approaching 90% for background sounds and nearly 100% for vocalizations (Fig. 7). Interestingly, the performance for identifying vocalizations improves with increasing sound duration while the performance for background sounds remains constant. This behavior occurs regardless of whether temporal, spectral or spectro-temporal correlations are used (Fig. 7, A-C). Since background sounds are more stationary, their statistics can be assessed quickly in this relatively simple task by the classifier. Vocalizations, on the other hand, are nonstationary over longer time-scales and have epochs of silence that may require the classifier to accumulate evidence over longer time. Additionally, the classification accuracy of background sounds is comparable (~90%) for spectral and temporal features. Except for the shortest sound duration used (100 ms), vocalization sounds are more accurately classified with spectral correlations compared to temporal correlations (p<0.01, t-test with Bonferroni correction), and the spectral correlations alone appear to account for most of the performance of the spectro-temporal classifier.

**Figure 7:**
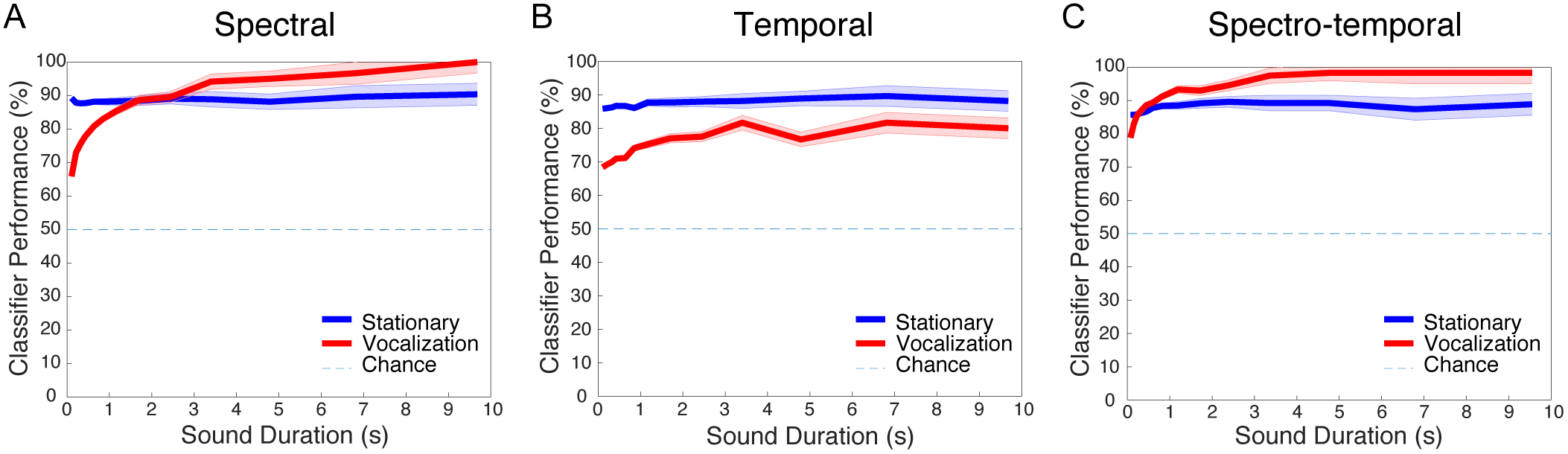
Classification performance in a two-category identification task distinguishing vocalization from background sound categories. For all three classifiers, the overall performance is consistently high and improved with increasing sound duration. Vocalization classification performance is highest for the (C) spectro-temporal classifier and shows a nearly identical trend for the (A) spectral classifier. The performance of the temporal classifier, however, is 20 % lower. For background sounds, classification performance did not improve over time and is consistently high (~90%) for all three classifiers.

## DISCUSSION

Here we have demonstrated that natural sounds can have highly structured spectro-temporal correlations and that this acoustic structure induces highly structured correlations between neural ensembles in the auditory midbrain. Stimulus-driven correlation structure, in both time and frequency (or space), is highly informative and conveys information about the sound identity, with evidence accumulation times on the order of a few seconds. Spectral and temporal correlations both contain information about sound categories and, nonstationary correlation structure conveys information beyond the corresponding time-average sound correlations. Altogether, these results demonstrate how time-varying correlations between modulations of sounds and the resulting correlations in frequency organized neural ensembles have the capacity to contribute to sound recognition.

### Using Sound Driven Correlation Statistics for Recognition

Previous work on how populations of neurons encode stimuli has emphasized the distinct roles of noise-driven and stimulus-driven correlations between neurons in limiting the information capacity of neural ensembles. Noise correlations, which are the result of coordinated firing not directly related to the stimulus structure, are thought to limit the encoding of sensory information ^19,20^. At the same time, theories of efficient coding suggest that neural ensembles should minimize the amount of stimulus-driven correlated firing to limit redundancies in the neural code ^10,11,18^. Here we find that stimulus-driven correlations between neural ensembles in the IC are highly structured. In contrast, the noise correlations in our neural data are largely unstructured. They are localized primarily to nearby recording channels as well as to brief time-epochs, and they do not vary systematically with the sound (Fig S2.1-2.3 and S3). These finding thus provide an alternative and, perhaps, underappreciated viewpoint: correlations in neuron ensembles can convey critical information about the stimulus itself. Rather than redundant structure that should be removed ^11^, sound correlation statistics can be viewed as high-level acoustic features or cues that are highly informative and which, as demonstrated, drive correlated activity in mid-level auditory structures.

Indeed, recent studies on sound textures confirm that correlation structure is a critical cue required to create realistic impressions of sounds ^2^. And there is growing evidence from neural recordings that neural correlations can convey information about stimuli or behavior during decision making ^30-32^. Our findings extend these views, by demonstrating that neural ensembles in a mid-level auditory structure can directly signal sound correlation structure and that sound categories have unique correlation statistics that may promote sound categorization.

### Biological Plausibility

While our results demonstrate that neural ensemble correlations in the auditory midbrain are highly structured in both the spectral and temporal domains, how and if such information is used by higher brain regions needs further exploration. One possible mechanism for representing correlations between neurons has been previously considered for pitch detection: the frequency coincidence detection network ^22,23^. The key proposal of this network is that neurons encoding different frequencies project onto the same downstream neurons that then detect coincident frequencies. Given that anatomical connections within and beyond the IC can span a broad range of frequencies, the central auditory system anatomy allows for such a possibility. Many projection neurons in the IC have collaterals that extend across multiple frequency-band lamina and send their outputs to auditory thalamus ^33^. Connections between thalamus and auditory cortex as well as intra-cortical connections, although frequency specific, can also extend across several octaves ^34,35^. This pattern of connectivity has the capacity to integrate information across frequency channels and may subserve a coincidence-like operation between inputs of different frequency. This coincidence detection would allow downstream neurons to directly compute spectral correlations, not just for pitch detection but, for sound categorization as well.

### Contribution of Spectral and Temporal Correlations

A key question brought up the model and neural data is the extent to which spectral and temporal correlations individually contribute to sound recognition. The auditory model suggests that both can contribute roughly equally for the sound categories tested, and performance is only marginally better when the two dimensions are combined. On the other hand, although the neural classification performance favors spectral (i.e. spatial) correlations over temporal, both sets of features have discriminative power. However, when the tonotpic organization is removed so that purely temporal correlations are used for the neural data, the temporal classifier performance is substantially reduced. This reduced performance is not observed in the model, since the temporal classifier used is specifically designed to isolate strictly temporal cues (see Materials and Methods). The observed difference between model and neural data may partly reflect differences in the sound paradigm and classifier used (single sound identification versus category identification). It is also possible that such differences are partly attributed to neural mechanisms not captured by the model, including spectro-temporal nonlinearities ^1^ or adaptation ^36^ at the level of the IC. As such, more extensive studies are necessary to parse out differences between the two domains.

Insofar as neural mechanisms for computing the sound correlation statistics, spectral models have broader biological support. Spectral correlations could be computed using simple coincidence detection, where two sound modulated inputs tuned to distinct frequencies converge, multiplicatively on downstream neurons. This type of spectral convergence is widespread in auditory system anatomy and the required multiplicative interactions have been previously described ^37,38^. On the other hand, computing temporal correlations requires coincidence detection of activity occurring at different times. Delay lines, differences in integration times, or feedback loops could all subserve these computations (with increasing temporal differences), but these phenomena are more speculative and lack strong anatomical or physiologic evidence.

### Resolution and Integration Time-Scales for Feature Analysis and Inference

Sound categorization performance for both the neural data and cochleogram model correlations improve over the course of 1-2s and depended strongly on sound duration, similar to human listeners ^39^. This brings up the question of whether the previously observed perceptual integration times in human observers should be attributed to a central neural integrator which averages sound statistics with this relatively long time-constant of a few seconds. Rather than computing average statistics about a sound over long time-scales, its plausible that sound statistics themselves are integrated and estimated at relatively short time scales analogous to the optimal integration resolution of our model (~150 ms). The long time-scales of a few seconds required to make perceptual decisions, may instead reflect a statistical evidence accumulation process, as previously proposed for cortical areas involved in decision making ^40^.

In modeling correlations in natural sounds, we find that the times required to accumulate statistical information about sound categories are roughly an order of magnitude larger than the optimal temporal resolution for calculating the correlations (~100-150 ms). This is consistent with a temporal resolution-integration paradox previously observed for neural discrimination of sounds ^41^. However, these time-scales are substantially longer than those previously reported for neural discrimination of sounds in auditory cortex. Previous studies identified an optimal temporal resolution of ~10 ms and an integration times of ~500 ms for discriminating pairs of sounds in auditory cortex ^41,42^. It is likely that such differences in temporal resolution and integration times largely reflect differences in the encoded features for our categorization paradigm and differences in how statistical evidence for the task is accumulated. The timescales for auditory cortex in these previous studies were optimized for the discrimination of pairs of sounds based on spike train distance measures at a single neuron level. By comparison, here we use high-order correlation statistics of either frequency-tuned cochlear channels or neuron ensembles as the primary feature for categorizing many more sounds.

### Conclusion

We have shown how the spectro-temporal correlation structure of natural sounds shows reliable differences between categories. For nonstationary sounds, this structure fluctuates on relatively fast (~150 ms) timescales even within a single sound. Surprisingly, this acoustic structure is reflected in the correlation structure of neural ensembles and can be used for accurate categorization with evidence accumulation time-scales of a few seconds. Together, our results from neural data and sound statistics suggest that spectro-temporal correlations in the auditory system may play an important role, not just in pitch perception or localization, but in sound recognition more generally.

## MATERIALS AND METHODS

### Animal Experimental Procedures

Animals are handled according to approved procedures by the University of Connecticut Animal Care and Use Committee and in accordance with National Institutes of Health and the American Veterinary Medical Association guidelines. Multi-channel neural recordings are performed in the auditory midbrain (inferior colliculus) of unanesthetized female Dutch-Belted rabbits (*N* = 2; 1.5-2.5 kg). Rabbits are chosen for these experiments since their hearing range is comparable to that of humans, and they sit still for extended periods of time which enables us to record from different brain locations daily over a period of several months.

### Surgery

All surgical procedures are performed using aseptic techniques. Surgical procedures are carried out in two phases with a recovery and acclimation period between procedures. For both procedures, rabbits are initially sedated with acepromazine and a surgical state of anesthesia achieved via delivery of isoflurane (1-3%) and oxygen (~2 liters/min). In the first procedure, the skin and muscle overlying the dorsal surface of the skull are retracted exposing the sagittal suture between bregma and posterior to lambda. Stainless steel screws (0-80) and dental acrylic are used to affixed a brass restraint bar oriented rostro-caudally and to the left of the sagittal suture. Dental acrylic is then used to form a dam on the exposed skull on the right hemisphere between lambda and the interparietal bone. Next, custom fitted earmolds are fabricated for each ear. A small cotton ball is inserted to block the external auditory meatus, and a medical grade polyelastomeric impression compound poured into the ear canal. After ~5 minutes, the hardened impression compound is removed. The ear impression mold is subsequently used to build a cast from which custom ear molds are fabricated.

Following the first surgery and a five-day recovery period, the animal is acclimated over a period of 1-2 weeks to sit still with the head restrained. During this period, the animal is also gradually exposed to sounds through the custom fitter ear molds. Once the animal is capable of sitting still during sound delivery, the second surgical procedure is performed. The animal is again anesthetized as described above, and an opening (~4 × 4 mm) is made on the right hemisphere within the dental acrylic dam and centered approximately 12-13 mm posterior to lambda. At this point, the exposed brain area is sterilized and medical grade polyelastomeric compound is poured into the acrylic dam to seal the exposed region.

### Sound Delivery and Calibration

Sounds are delivered to both ears via dynamic speakers (Beyer Dynamic DT 770 drivers) in a custom housing and custom fitted ear molds obtained as described above. The molds are fitted with a sound delivery tube (2.75 mm inner diameter) that is connected to the dynamic speaker housing forming a closed audio system. Calibration consisted of delivering a 10-sec long chirp signal at 98kHz sampling rate via TDT RX6 and measuring the audio signal with a B&K calibration microphone and probe tube placed ~5 mm from the tympanum. The measured signal is used to derive the sound system impulse response (via Wiener filter approach; combined speaker driver and tube) and an inverse filter finite impulse response is then derived. Subsequently, all sounds delivered to the animal are passed via the inverse filter which is implemented in real time using a TDT RX6 at 98kHz sampling rate. The sound delivery system has a flat transfer function and linear phase between 0.1 – 30kHz (flat to within ~ ±3 dB).

At each penetration site, we first delivered a pseudo random sequences of tone pips (50 ms duration, 5 ms cosine-squared ramp, 300 ms inter-tone interval) covering multiple frequencies (0.1-16 kHz) and sound pressure levels (5-85 dB SPL). These tone-pip sequences are used to measure frequency response areas which allow us to estimate the frequency selectivity of each recording site (different channels).

Next, we delivered a sequence of five environmental noises with distinct structural properties to determine whether the ensemble activity of the auditory midbrain reflected the sound frequency dependent correlation statistics present in the sounds. Each sound is 3 s duration and is delivered in a block randomized fashion with a 100 ms inter-stimulus interval between sounds. To avoid broadband transients, the sounds have b-spline onset and offset ramps (20 ms rise-decay time) and are delivered at an RMS sound pressure level of (70 dB SPL). The sounds included: running water, a crackling fire, speech babble, a bird chorus, and a rattling snake sound. These sounds each contain unique time-frequency correlation statistics allowing us to test and quantify whether such statistics are potentially encoded by the auditory midbrain and ultimately represented in the neural ensemble activity. For instance, the water sounds have minimal across-frequency channel correlation since the air bubbles and droplets responsible for this sound are relatively narrow band and occur randomly in time, thus activating frequency channels independently ^7^. Sounds such as crackling fire, by comparison, have strong frequency dependent correlations due to crackling embers which produced brief impulsive “pops” that span multiple frequency channels simultaneously. The temporal correlations of these five sounds are also quite varied. For instance, the water sound has a very brief impulsive correlation structure lasting just a few milliseconds whereas the bird chorus and speech babble have a broader and slower temporal correlation function. The rattling snake sound, by comparison, had strong periodic correlations at ~20 Hz.

### Electrophysiology

Sixteen channel acute neural recording silicon probes (Neuronexus 10 mm probe; 16- linear spaced *recording* sites with 100 um separation; site impedance ~1-3 *M*Ω) are used to record neural activity from the inferior colliculus of unanesthetized rabbits. We recorded neural data from N=13 *penetration* sites in the inferior colliculus of two rabbits (N=4 and N=9). Since there are 16 recording channels for each penetration site, data is obtained from a total of 16×13=208 recording sites within IC. Prior to recording, the polyelastomeric compound is removed from the craniotomy and lidocaine is applied topically to the exposed cortical tissue. The area is then flushed with sterile saline and the acute recording probe inserted at ~12-14 mm posterior to lambda. If necessary, a sterile hypodermic needle is used to nick the dura to allow the electrode to penetrate the neural tissue. An LS6000 microdrive (Burleigh EXFO) is used to insert the neural probe to a depth of ~7.5-9.5 mm relative to the cortical surface where, at this penetration depth, most or all of the recording electrodes are situated in the IC and had clear responses to brief bursts of broadband noise or tones. Neural activity is acquired continuously at sampling rate of 12 kHz using a PZ2 preamplifier and RZ2 real time processor (TDT, Alchua, Florida).

The sampled extracellularly recorded neural signals are analyzed offline using an analog representation of multi-unit activity (analog multi-unit activity, aMUA)^43^. From each of the recorded neural traces aMUA is measured by first extracting the envelope of the recorded voltage signal within the prominent frequency band occupied by action potentials spanning frequencies 325 and 3000 Hz (b-spline filter, 125 Hz transition width, 60 dB attenuation) ^44^. The bandpass filtered voltage signal is next full-wave rectified and low-pass filtered with a cutoff frequency of 475 Hz (b-spline filter, transition width of 125 Hz and stopband attenuation of 60 dB), since neurons in the auditory midbrain typically don’t phase lock to envelopes beyond ~500 Hz ^45^. The resulting envelope signal is next down-sampled to 2 kHz. Such neural envelope signals captures the synchronized activity and the changing dynamics of the local neural population with each recording array in both time and frequency domains ^43^. For each recording channel an analog raster is generated which consists of the aMUA response over time and across trials. Each recording had 5 sound conditions where each sound had at least 18 trials (range 18 to 39) with a 3 s duration for each trial.

### Neural Ensemble Stimulus-Driven Correlation Matrix

For each recording penetration, we estimat the *stimulus-driven* correlations of the neural ensemble across the 16-recording channels directly from the measured aMUA signals. The procedure consists of a modified windowed short-term correlation ^27^ analysis between recording sites in which the correlations are “shuffled” across response trials ^18,46^. The windowed correlation approach allows us to localize the correlation function in time, whereas the shuffling procedure is used to remove neural variability or noise from the correlation measurements. Thus, the proposed shuffled windowed correlation allows us to isolate stimulus-driven correlation between recording sites independently of noise driven correlations. The windowed shuffled cross-correlation between the *k*-th and *l*-th recording site is computed as the mean pairwise correlations between different trials of the aMUA envelope:

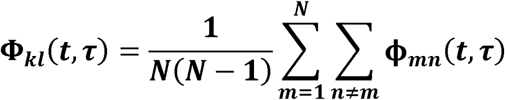

where *N* is the number of trials in each recording channel (*k* and *l)*. ϕ_*mn*_(*t, τ*) is the windowed cross-correlation between the *m-*th and *n-*th response trials in the two channels respectively, where *t* is time and *τ* is the cross-correlation delay

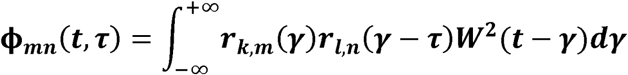

where γ is the time integration variable. Here *r*_*k,m*_(*t*) is the *m*-th response trial (mean removed) from channel *k* and *r*_*l,n*_(*t*) is the *n*-th response trial (mean removed) from channel *l*. *W*(*γ*)is unit amplitude square window centered about γ=0 of duration *T* sec (range = 62.5 – 1000 ms) which is used to localize the measured signal correlations around the vicinity of the designated time-point (*t*). Note that in this formulation, correlations between recording channels (*k* and *l*) are computed for different response trials (*m* and *n*). The above is implemented using a fast-shuffled correlation algorithm according to ^46^:

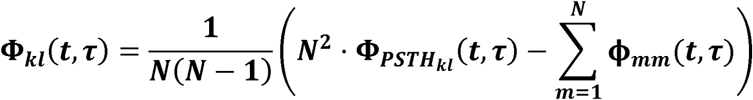

where Φ*_PSTH_kl__(t, τ)* is the windowed cross-correlation function for the post-stimulus time histograms (PSTHs) between channel *k* and *l*:

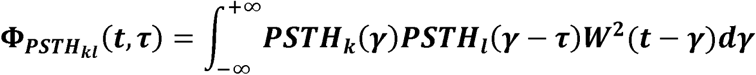

where the PSTH for the *k*-th and *l*-th channels are

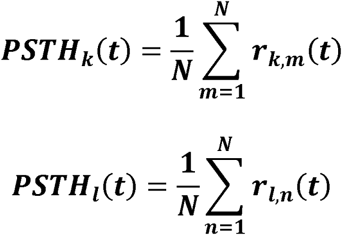

As previously shown ^46^, this fast-shuffled correlation algorithm resulted in a marked reduction in the computational time (*N*+1 correlations compared with *N*(*N*–1) for each pair of recording channels. This speedup in the computational time is necessary to bootstrap the data during the model generation and validation applied subsequently (See *Neural Ensemble Classifier*).

To remove the influence of the ressponse power on the correlation measurements, the short-term correlation is normalized as a correlation coefficient. This requires that we measured the localized short-term signal variance at each time and delay sample according to

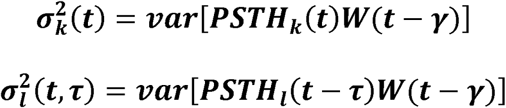

The channel-covariance is then obtained as

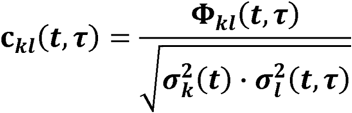

Like a correlation coefficient, this population ensemble correlation function is bounded between −1 and 1.

### Neural Ensemble Classifier

We use a cross-validated neural ensemble classifier to assess whether spatial, temporal, and/or spatio-temporal correlations of the neural ensemble in IC could be used to recognize sounds. Although technically, the correlations are measured between the neural activity across spatially separated electrode channels (*k* vs. *l*), the electrode channels with our recording paradigm are frequency ordered (Fig. 1A). The neural ensemble activity thus reflects spectral correlations between frequency channels, and, in what follows, we describe spatial correlations between recording channels as spectral correlations.

The neural classifier is implemented using a cross validation approach in which half of the data is used for model generation and the second half for model validation. For each and the second half as the validation data. The first and second half of the data are then swapped and the procedure repeated using the first half of the data for validation and the second half for model generation. The model consisted of the time average correlation function at zero-lag for each of the five sounds *C_kl,s_* (0), where *s* indicates the sound. Note that, since the model correlations are averaged across time, *t* is no longer a variable in the model correlation. The model classification performance is tested and validated by iteratively implementing a minimum Euclidean distance classifier across different validation data segments according to:

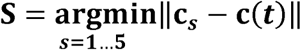

where *s=*1…5 are the five sounds tested, S is the classified sound, *c*(*t*)is a vector of features obtained from the spectro-temporal correlation function for a particular time segment of the validation data (at time *t*) and c_s_ is a vector containing the average correlation feature vector for sound *s* from the model generation data. The center location of each correlation measurement, *t*, is randomly varied by randomly sampling 100 distinct time points (*t*). The analysis is repeated for window sizes ranging between 62.5 and 1000 ms in ½ octave steps. The overall classifier performance is the average across all five sounds and across the 100 randomly selected validation segments.

To evaluate the contribution of temporal and spectral correlations for neural classification performance, we implement the neural classifier either using purely temporal, purely spectral, or joint spectro-temporal correlations. The purely spectral classifier only considers correlations at zero lag, *c_spec_*(*t*)=*c_kl_*(*t*,0), as the primary features (no time lag between different frequency channels). Note that *c_kl_*(*t*,0)contains strictly frequency dependent information, since the recorded neural channels are tonotopically ordered and delays are removed. For the temporal classifier, we consider the correlations along the diagonal, c_*temp*_(*t*) =c_*kk*_(*t,τ*), for delays extending between τ = −100 to 100 ms as the primary features. Next, we combined the spectral and temporal correlations to implemented the joint spectro-temporal classifier as follows:

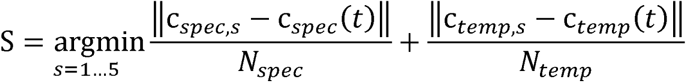

where *N_spec_* and *N_temp_* are the number of elements contained in the spectral and temporal feature vectors. By normalizing the spectral and temporal distance measures by the number of elements in each vector we ensure that neither the spectral and temporal correlations dominate the categorization.

Finally, we implement a purely temporal correlation classifier that lacks frequency organization. Note that for the temporal correlation classifier, the features consist of the envelope autocorrelations taken across all possible frequency channels (*c_kk_*(*t,τ*)), an can convey tonotopic information through the identity of the recording channel *k*. In order to isolate purely temporal correlation cues, we remove the tonotopic information from the temporal correlation signal by randomly reordering the frequency channels during the classification step.

#### Sound Database for Auditory Model and Classifier

Sounds representing 13 acoustic categories are obtained from a variety of digital sound sources. 195 sounds from 13 different acoustic categories are used to build distributions of dynamic, spectro-temporal correlation statistics of natural/man-made sounds. Sound segments are chosen so that they have minimal background noise and are drawn from 3 broad classes: vocalizations, environmental sounds, and man-made noises. Vocalizations include 1) Single bird songs (various species), 2) Cat meowing (single or multiple cats), 3) Dog barking (single or multiple dogs) and 4) Human speech (male speaker). Environmental sounds include 5) Bird chorus (various species), 6) Speech babble (in various environments e.g. bars, super markets, squares), 7) Fire, 8) Thunder rain, 9) Flowing water (rivers and streams), 10) Wave (ocean/lake waves), 11) Wind. Finally, man-made noises consist of 12) Bell (church or tower bells) and 13) Automobile engines (different vehicles). Each category contains 15 sounds, each 10 s long, sampled at *F_s_* 44.1kHz (see Table S1 for sources and full list of tracks used).

#### Auditory Model

Sounds are analyzed through a cochlear filter bank model of the auditory periphery that decomposes the sound using frequency organized cochlear filters. The cochlear filter banks consists of tonotopically arranged gamma-tone filters (Irino & Patterson, 1996). These filters have a sharp high frequency cutoff and shallow low frequency tails that resemble the tuning functions of auditory nerve fibers. The *k*-th gamma-tone filter has an impulse response function:

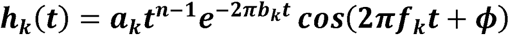

where *k* is the filter channel, *t* denotes time, *b_k_*, and *f_k_* denote the filter bandwidth and center frequency. The filter gain coefficient *a_k_* is chosen so that the filter passband gain is 1; filter order *n* and filter phase *ϕ* are 3 and 0, respectively. Filter bandwidths are chosen to follow perceptually derived critical bandwidths 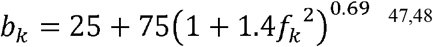. We use *L*=58 frequency channels with center frequencies *f_k_* ranging from 100 Hz to 16 KHz in 1/8 octave steps. In the first stage of processing, the sound *s*(*t*) is passed through the cochlear filterbank model:

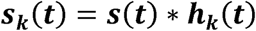

where * represents the convolution operator. The outputs of the filter bank are next passed through a nonlinear envelope extraction stage, which models the characteristics of the hair cell. We first compute the magnitude of the analytic signal (Cohen,1995):

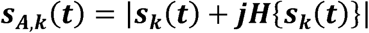

where H{·}is the Hilbert transform operator and 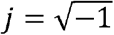. Then the temporal envelope for each channel is obtained by convolving the rectified analytic signal magnitude with a b-spline low pass filter with cutoff frequency of 500 Hz (transition bandwidth of 125 Hz and stopband attenuation of 60 dB):

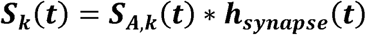

which models the low-pass filtering of the hair cell synapse (*h*_synapse_(t)). The lowpass filtered envelopes are then down-sampled to 1 kHz for modeling. Here we refer to the time-varying env e,lopes of the cochlear filters as the c,ochlear spectrogram and use the notation *S*(*t*,*f_k_*)=*S*_k_(*t*). The cochlear spectrograms, *S*(*t*,*f_k_*), of all sound segments are normalized to have zero mean and unit variance.

#### Nonstationary Spectro-temporal Correlation Statistics

Since natural sounds are often nonstationary, we measure not just the long-term correlation, but the time-varying or “short-term” correlation statistics between the frequency organized channels in the cochlear model representation. This nonstationary representation is then used to quantify the contribution of the sound correlation statistics to sound categorization. The short-term correlation statistics that we use are similar to those for the neural data analysis except that we use the frequency channels from the cochlear spectrogram rather than the neural signals. The running short-term correlation function Φ between the cochlear spectrogram channel *k* and *l* is computed according to ^27^:

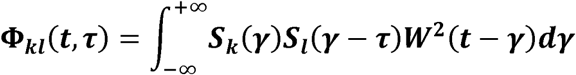

where *W*^2^(γ) is a sliding window function that determines the temporal resolution of the correlation measurement, *t* is the time, τ is the cross-correlation delay. Here *W*^2^(γ) is a Kaiser window (*β* = 3.4) where the overall window resolution (corresponding to two standard deviations of the Kaiser window width; varied between 25 to 566 ms in ½ octave steps) is varied to quantify the effects of the correlation temporal resolution on categorization performance. The range of permissible cross correlation delays (–τ_*W*_ to + τ_*W*_) correspond to half the window size and is thus varied between −12.5 to +12.5 for the highest resolution to −283 to +283 ms for the coarsest resolution. Conceptually, at each time point, the short-term correlation performs a correlation between the locally windowed envelope signals *S_k_*(*t*)*W*(*t*–*γ*) and *S_l_*(*t*–*τ*)*W*(*t*–*γ*) to estimate the localized correlation statistics of the cochlear envelopes.

To remove the influence of spectral power on the correlation measurements, the short-term correlations are normalized

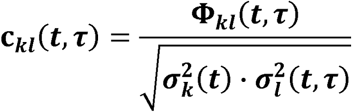

where

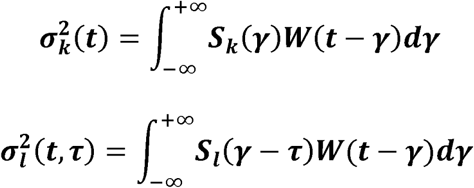

are the time-varying and delay varying (for channel *l*) power *k*-th and *l*-th spectrogram channels. Again, as with a Pearson correlation coefficient, this short-term spectro-temporal correlation is bounded between −1 and 1.

To assess the contribution of temporal and spectral correlations on the sound categorization performance, we perform a secondary analysis where the spectro-temporal correlation function is decomposed into its purely spectral or temporal components, following a similar framework as for the neural data analysis. To evaluate the spectral correlations, we consider only the correlations at *τ* = 0 (no time lag between different frequency channels). For temporal correlations, *τ* ranges from 0 up to half of the Kaiser window length. Only auto-correlation functions or correlation functions of a channel vs. itself are considered in temporal correlation analysis although all possible frequency channels are involved in spectral correlation analysis. Correlations between different frequency channels computed at zero time-lags are referred to as purely spectral correlation components whereas the auto-correlations computed at different time lags for each frequency channel are referred to as temporal correlation throughout.

#### Stationarity and Ensemble Diversity Indices

Given that sounds in our database are quite varied, ranging from isolated vocalizations to environmental sounds consisting of superposition of many individual acoustic events, we seek to characterize the overall degree of stationarity in the short-term correlation statistics for each of the ensemble. Furthermore, since sound recordings are obtained from different sources and animal species (e.g., for vocalizations) all of which could influence the overall category statistics we also seek to quantify the overall diversity of the short-term correlation statistics of each ensemble. When considering sound categorization, we might expect that stationary sounds with minimal diversity across an ensemble would be most easily recognized.

For each sound, the sampled short-term spectro-temporal correlation c_*kl*_(t,τ) (computed using τ_W_=100 ms) is rearranged and expressed as a time-dependent vector function c̄(*t*) with dimensions *M L*^2^ at each time point, where *M*=99 is the number of time lags used for the short-term correlation and *L* is the number of frequency channels. The stationarity index of each sound is defined and calculated as

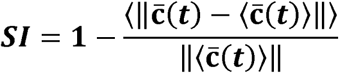

Where ||·||is the vector norm and 〈·〉 is a time-average. Conceptually, the SI corresponds to the time-average variance of the short-term correlation normalized by the total power of the time-averaged short-term correlation. As such, it measures the average normalized variability across time and is bounded between 0 and 1, where 1 indicates that the moment-to-moment variance of the short-term correlation is 0 thus indicating a high degree of stationarity. By comparison, when the moment-to-moment variance is high the index approaches 0 indicating a highly nonstationary short-term correlation function.

We next define and measure the category diversity index (*CDI*), which is designed to measure the degree of homogeneity or heterogeneity in the short-term spectro-temporal correlation functions for each of the 13 sound categories studied. To do so, we first compute the time-average correlation function for each sound in a given ensemble, c̄_*n*_ = 〈c̄_n_(*t*)〉, where *n*=1…15 is an index representing the sounds for each sound category. The *CDI* is defined and computed as

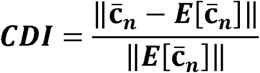

where *E*[·] is the expectation operator taken across the sound ensemble (equivalent to an average across sounds, *n)*. Conceptually, the *CDI* corresponds to the variance of the time-average short-term correlation taken across the ensemble of sounds normalized by the power (norm) of the time and ensemble average short-term correlation.

For a particular sound category, a *CDI* near zero is indicative of low diversity (homogeneity) such that the short-term spectro-temporal correlation functions of that ensemble are quite similar from sound-to-sound and thus closely resemble the average ensemble correlations. By comparison, an *CDI* of 1 indicates a high degree of heterogeneity (high diversity) so that the short-term spectro-temporal correlations are quite different from sound-to-sound.

#### Dimensionality Reduction and Distribution Model

To reduce the dimensionality of the categorization problem, we use principal component analysis (PCA). For spectral correlation statistics, the entries of the correlation matrix at zero time-lag (c_*kl*_(*t*,0)) are considered as features while time points are used as observations or trials. For temporal correlation statistics, the correlations at different time lags within single frequency channels (c_*kk*_(*t*,τ)) are considered features. Both time points and different frequency channels are treated as observations so that temporal information is not specific to any particular frequency channel. For further analysis we use only the highest ranked principal components that explain 90% of the variability in the data (26 PCs for spectral; 8 for temporal; 87 for spectro-temporal).

Using the low-dimensional representations of the spectral, temporal, or spectro-temporal correlations, we model the distributions of principal components for each sound category with a Gaussian mixture model (GMM). For each sound category *i* we learn a multivariate probability distribution:

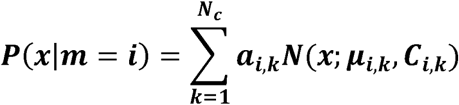

where *x* is the low-dimensional vector of PCA scores, *m* are the sound categories, *p*(*x*|*m*)is the multivariate PDF of sound features for category *m*, *a*_k_≥0 is the weight of the *k-*th mixture component, *N*(*x*;*μ_k_*,*C_k_*)is the PDF of a multivariate normal distributions with mean *μ_k_* and covariance *C*_*k*_, and *N*_c_ is the number of Gaussian components used for modeling. To avoid ill-conditioned covariance matrics, we constrain *C_*k*_* to be diagonal.

In order to find the optimum number of Gaussian components (*N_c_*) required to model the data, we compute cross-validated likelihoods for different numbers of Gaussian mixtures (*N*_c_=1 to 20). The optimal *N_c_* values are 5, 8 and 13 for temporal, spectral, and spectro-temporal, respectively.

#### Bayesian Classifier

Given the mixture model for each sound, we then use a Bayesian classifier for sound identification. The features fed to the classifier consist of the principal component scores from either the sound’s short-term spectral, temporal or spectro-temporal correlation. Since we are feature vectors x = [*x*_1_,*x*_2_,…,*x_N_*]consisting of principal component scores obtained from successive, windowed segments of sounds at different time samples (*t*_1_…*N*) selected so that adjacent sound segments do not overlap. We then evaluate the posterior probability of each sound segment under the different mixture models. The most probable case is chosen according to the maximum a posteriori (MAP) decision rule:

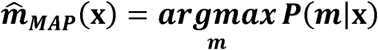

In practice, we find the MAP category by using Bayes rule and maximizing the log-likelihood. We assume that the categories are equi-probable a priori and that the features at each time sample are conditionally independent. Thus, the MAP category is obtained by maximizing:

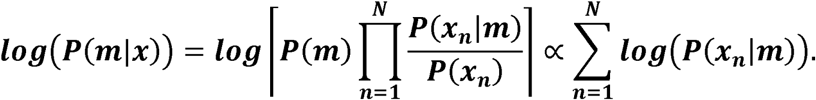

#### Cross Validation

To avoid over-fitting, we use a leave-one-out cross-validation, where all sounds are used to build the model distributions except for one sound which is used for testing (Larose & Larose, 2015). Because there is only one sound validated, each validation iteration produces a 0 % or 100 % correct classification rate. The procedure is repeated iteratively over all sounds and the average performance is obtained as the average classification rate across all iterations. The total sound duration used for the validation is varied by selecting *N* consecutive time window segments as described above from each sound under the test to be categorized; the selected *N*-segments start from the very beginning of the sound up to the end of the sound. Values of *N* are varied in 1/2 octave steps starting with *N*=1 up to the maximum value allowed by the sound duration.

#### Optimal Temporal Resolution and Integration Time for Categorizing Sounds

As previously demonstrated for auditory neurons an optimal temporal resolution can be identified for neural discrimination of natural sounds, and neural discrimination performance improves with the increasing sound duration ^41,42^. For this reason, we seek to identify both the optimal temporal resolution that maximizes categorization performance as well as the integration time of the sound classifier. The temporal resolution of the correlation signals is varied by changing the sliding window temporal resolution, τ_*W*_, between 25-566 ms in ½ octave steps. Classification performance curves vary with τ_*W*_, exhibiting concave behavior with a clear maximum that is used to identify the optimal window time-constant. The classifier performance also increased in an approximately exponential fashion with the overall sound duration. The classifier performance also increases with the overall sound duration. The classifier integration rise-time, τ_*c*_, is defined as the amount of time required to achieve 90% of the asymptotic performance measured at 10 sec duration.

